# Deforestation and bird habitat loss in Colombia

**DOI:** 10.1101/2020.05.30.125849

**Authors:** Pablo Jose Negret, Martine Maron, Richard A. Fuller, Hugh P. Possingham, James E.M. Watson, Jeremy S. Simmonds

## Abstract

Tropical forests harbor most of the planet’s terrestrial biodiversity, and their loss means destruction of habitat for many species. Tropical deforestation continues at high rates in many regions, but it is often reported only in terms of area lost or its impacts on high-profile threatened species. We estimated the impact of both past and projected future deforestation on habitat extent for the entire assemblage of forest-dependent birds across Colombia, the country with more bird species than any other. Of the 550 forest-dependent species analysed, Almost all (n=536; 96.5%) had lost habitat, and 18% had lost at least half of their habitat by 2015. We used the recently developed Loss Index (LI) to capture the severity of habitat loss for the forest bird assemblage, discovering that the current LI for Colombia is 35, which means 35% of bird species have lost at least 35% of their habitat. The national LI for Colombia is projected to rise to 43 by 2040 if recent forest loss trends continue. There were large regional differences; Caribe had an LI of 82 while for the Pacific it was 14. A threat assessment for the regionally endemic species in the country showed that 12 (30%) of the species that are projected to lose 50% or more of their historical habitat by 2040 are not currently classified as threatened by the IUCN, suggesting that there are many species that are not listed but that face an imminent extinction threat from habitat loss. This extensive habitat depletion affecting entire species assemblages has significant implications for tropical forest ecosystems, and risks eroding ecosystem function and ecosystem service provision.

## Introduction

Bird species have been experiencing rapid population declines worldwide (Inger et al. 2015; Lindenmayer et al. 2018; Rosenberg et al. 2019), with habitat loss being one of the principal drivers (Owens & Bennett 2000; Ford et al. 2009; Calvert et al. 2013). These population declines and the local extirpation of species have been linked to a deterioration in the function of ecosystems and the provision of ecosystem services, such as pollination and insect control (Sekercioglu 2006; Gaston & Fuller 2008; Winfree et al. 2015). However, the impact of habitat loss for entire bird assemblages has been little studied in tropical ecosystems (Sekercioglu 2012), despite the fact that such ecosystems harbor most of the planet’s terrestrial biodiversity (Gibson et al. 2011). Moreover, there is clear evidence that species-rich assemblages such as those that characterise tropical forests play a critical role in ecosystem function and service provision (Sekercioglu 2012; Gaston et al. 2018).

In the last two decades more than 2.3 million square kilometres of tropical forest have been lost (Hansen et al. 2013; World Resources Institute 2018) and the destruction of these habitats threatens the survival of forest specialist species (Donald et al. 2018; Symes et al. 2018; Watling et al. 2020). As tropical forests continue to decline in extent (Malhi et al. 2014), habitat for a huge number of species is being lost (Ceballos et al. 2017; Lovejoy 2017). However, we lack a clear picture of what this loss means for both whole assemblages of species, and particular subsets of assemblages that are of conservation interest (e.g. endemic species, functional groups; Simmonds et al. 2019).

Here, we assess the implications of forest loss for bird assemblages in Colombia, the country with more bird species than any other (Ayerbe-Quiñones 2018), and where forest loss is particularly acute (Negret et al. 2019). We estimate the impact of past and projected deforestation on the extent of potential habitat for 550 forest-dependent bird species. Historical, recent, and projected future habitat loss for forest-dependent species was calculated across Colombia, as well as for each biotic region, for species-rich taxonomic groups of the broader assemblage (tanagers, ovenbirds, flycatchers and antbirds) and separately for 69 regionally endemic forest-dependent species (defined as having ≥80% of their global distribution in Colombia). In this way the impact of habitat loss was assessed for all the forest-dependent species in the assemblage.

## Methods

### Study area

Colombia, in north western South America, is a tropical country of 1,142,000 km^2^ with a wide topographic range. As a result, the country has extraordinarily high vertebrate diversity including >1900 species of birds (McMullan et al. 2010; Ayerbe-Quiñones 2018), 700 species of amphibians (Galeano et al. 2006) and 400 species of mammals (Ramirez-Chaves & Suárez-Castro 2014). Natural forest covered more than 55% of its continental area in 2018 (World Resources Institute 2018).

### Distribution ranges for forest-dependent species

We obtained distribution and ecological data for all native birds occurring in Colombia from BirdLife International using the latest available range maps (BirdLife International & Handbook of the Birds of the World 2018). These distribution maps are generated by experts based on their knowledge and the available data for each species (BirdLife International & Handbook of the Birds of the World 2018; IUCN Red List Technical Working Group 2019). We then filtered our list to include only forest-dependent bird species following the Donald et al. (2018) definition of forest-dependent species: those whose listed habitat as defined by the IUCN habitat classification scheme (https://www.iucnredlist.org/resources/habitat-classification-scheme) included only the level 1 classification “Forest & Woodland”. After filtering out non-forest and multi-habitat species, 550 forest-dependent species remained for analysis. We adhered to the BirdLife taxonomic treatment.

### Forest cover data

We used maps of forest cover in Colombia for four points in time (historical, 2000, 2015 and 2040) to determine the extent of suitable habitat inside each species’ range in each time period. For the distribution of forest cover in 2000 and 2015 we used the 1 km^2^ resolution forest cover maps generated by Negret et al. (2019). For the historical distribution of forest cover we used the map of historical cover of forest ecosystems in Colombia created by Etter et al. (2017). This map used Landsat images for the country from 1972 – 1977, a combination of different ecosystem maps (Etter 1998; Etter et al. 2006b) and information on the distribution of areas of historical change where deforestation for agricultural land uses have occurred, to define the potential distribution of the extent of forest cover if human intervention and transformation had not occurred (Etter et al. 2006b, 2017). The resolution of this forest cover map was 250 m^2^ and so we generated a 1 km^2^ grid covering Colombia to match the resolution of the forest cover map generated by Negret et al. (2019), and calculated the proportion of forest cover for each grid cell. We then defined grid cells with >30% forest cover as forest and those with <30% as non-forest based on the threshold used by the Colombian Institute of Hydrology, Meteorology and Environmental Studies – IDEAM (Galindo et al. 2014; Negret et al. 2019). In forest landscapes that retain less than 30% forest cover, bird species richness is markedly lower than those with greater cover (Ochoa-Quintero et al. 2015). Any pixel that was classified as non-forest using the historical forest cover layer was treated as no forest for all subsequent time slices.

### Forest cover change model

We simulated the spatial distribution of deforestation across Colombia in 2040, using a cellular automata model developed in Dinamica EGO with parameters that allocate deforestation on the basis of its empirical association with a set of predictor variables (Soares-Filho et al. 2002, 2013) including; proximity to roads, rivers, mining concessions, oil exploitation wells, distance to previous deforested areas, armed conflict intensity, distance to coca plantations, the presence of protected areas, slope and elevation (Negret et al. 2019). To do this, we assessed the association of the predictor variables with deforestation from 2000 to 2015 with the Bayesian weights of evidence method (Bonham-Carter 1994; Soares-Filho et al. 2013). Then we used the weights of evidence coefficients from the spatial determinants of forest change as inputs in a multi-stage process to model the spatial distribution of deforestation pressure in the country (Soares-Filho et al. 2002, 2013; Negret et al. 2019). The model used the weights of evidence coefficients, the 2015 forest cover map and the spatial distribution of the biophysical and anthropogenic variables to produce a spatial map of deforestation pressure. We then used an average annual deforestation rate calculated from the 2000 and 2015 forest cover maps and the deforestation pressure map to generate a forest cover projection for 2040 using Dinamica EGO software (Teixeira et al. 2009; Molin et al. 2017). Although the published model included the effect of armed conflict in the context previous to the signing of the peace deal with FARC, there has not been a cessation of conflict in the country (Parkins 2019), so the same model was used.

### Habitat loss across bird ranges

To estimate change in the extent of suitable habitat for each forest-dependent species at each of the four points in time (Historical, 2000, 2015 and 2040), we assumed all forest inside each species’ range was potential habitat. We quantified the total area of habitat for each species in each time period, the loss through time, and the proportion of the estimated habitat extent that the loss represented.

### Deforestation impact on species assemblages

We estimated the impact of deforestation up to 2015 for all the forest-dependent species in Colombia and for each biotic region. We also estimated the impact of deforestation for regionally-endemic species (defined as having ≥80% of their range in Colombia) and for the four most species-rich families (tanagers, ovenbirds, flycatchers and antbirds) to explore how forest loss has variously affected these bird groups. Finally, we assessed the projected impact of deforestation by 2040, for each biotic region, for the four species-rich taxonomic groups and for all regionally-endemic species. We used the Loss Index (LI) to describe potential natural habitat loss for the forest-dependent bird assemblages of Colombia (Simmonds et al. 2019). The LI is a metric where an LI of *x*, indicates that *x*% of species in an assemblage have each lost at least *x*% of their potential natural habitat (Simmonds et al. 2019).

### Threat assessment for regionally-endemic species

We performed a threat assessment for the regionally-endemic species (defined as having ≥80% of their range in Colombia) - for these species long-term survival is heavily dependent on their persistence in Colombia. We used habitat loss as a proxy for population decline assuming that the rate of habitat loss was directly proportional to the rate of population decline, such that a loss of 1% suitable habitat was equivalent to a 1% population decline (Symes et al. 2018). We then determined the proportional population decline until 2015 (historical loss), the decline to 2040 (projected loss), and the decline between 2000 and 2015 (recent loss) for each species. Using these data, we identified the species that had lost more than 20, 30, 50 and 80 percent of their population in the different timeframes (Historic, projected and recent loss) and contrasted this loss with the threat classification thresholds of population reduction from the IUCN Red List criterion A4 (IUCN Standards and Petitions Committee 2019); Critically Endangered (>80% reduction), Endangered (>50% reduction), Vulnerable (>30% reduction) and Near Threatened (>20% reduction). We are aware that these thresholds refer to change over 10 years or 3 generations for classification purposes, and use them here simply as a point of reference.

We also identified the species with area of occupancy of less than 3000, 2000, 500 and 10 km^2^ in 2015 and in 2040 and contrasted this with the threat classification thresholds for extent of area of occupancy from the IUCN Red List criterion B2 (IUCN Standards and Petitions Committee 2019); Critically Endangered (<10 km^2^), Endangered (<500% km^2^), Vulnerable (<2000% km^2^) and Near Threatened (<3000% km^2^). We defined the area of occupancy as the area inside each species’ range with forest habitat.

## Results

### National level

The vast majority of forest-dependent birds (n=536; 96.5%) in Colombia had lost suitable habitat by 2015 in comparison with their historical habitat extent (Fig. 1 & Supplementary Fig. 1). The average loss of habitat was 24% (± 20 SD) by 2000, 27% (± 20.6 SD) by 2015, and we projected an average of 38% (± 26 SD) loss of habitat by 2040. Only 17% (n=35) of the species that are projected to lose at least half their habitat in Colombia by 2040 (n=208) are currently classified as threatened by the IUCN.

**Figure 1.**
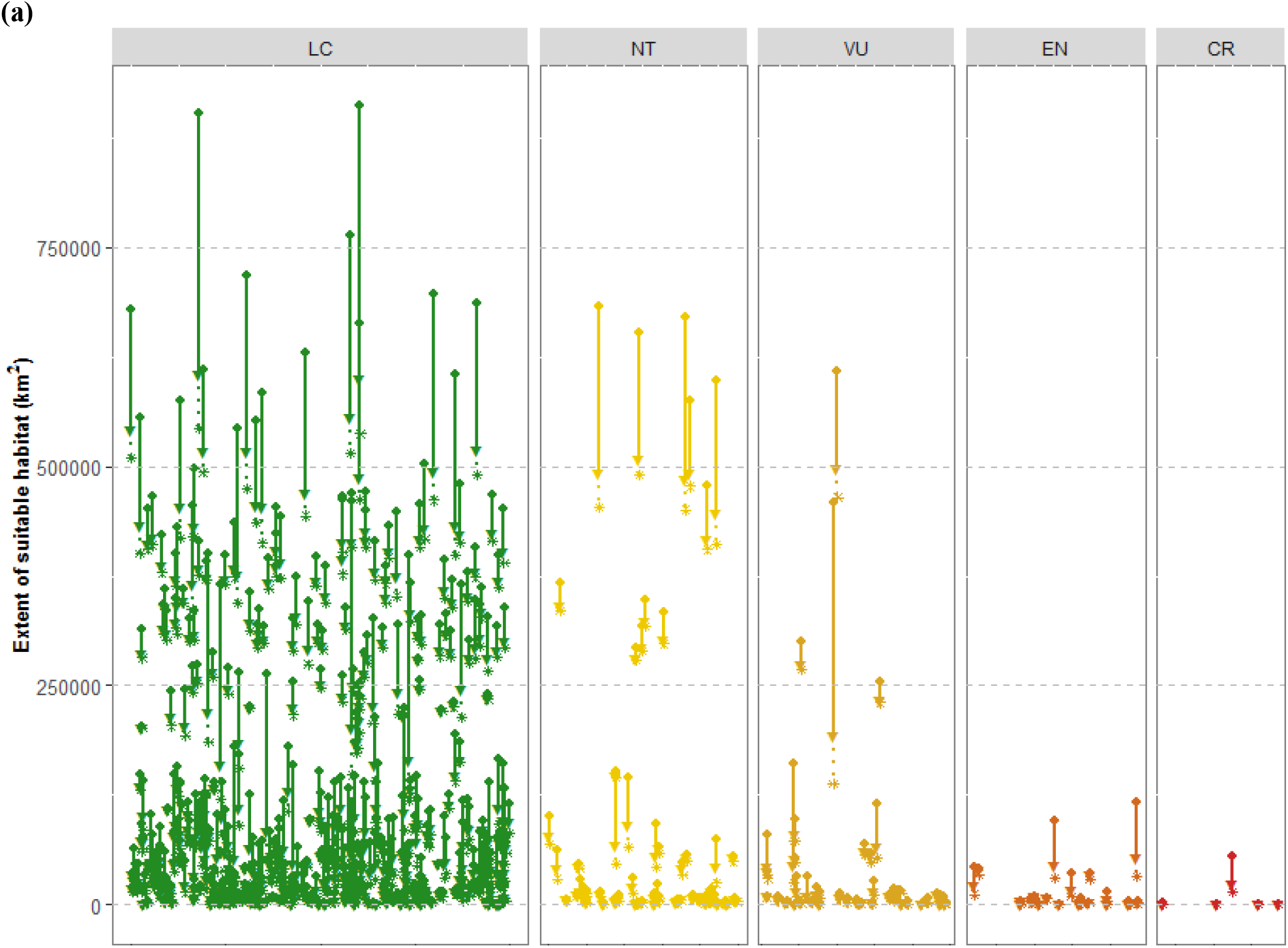
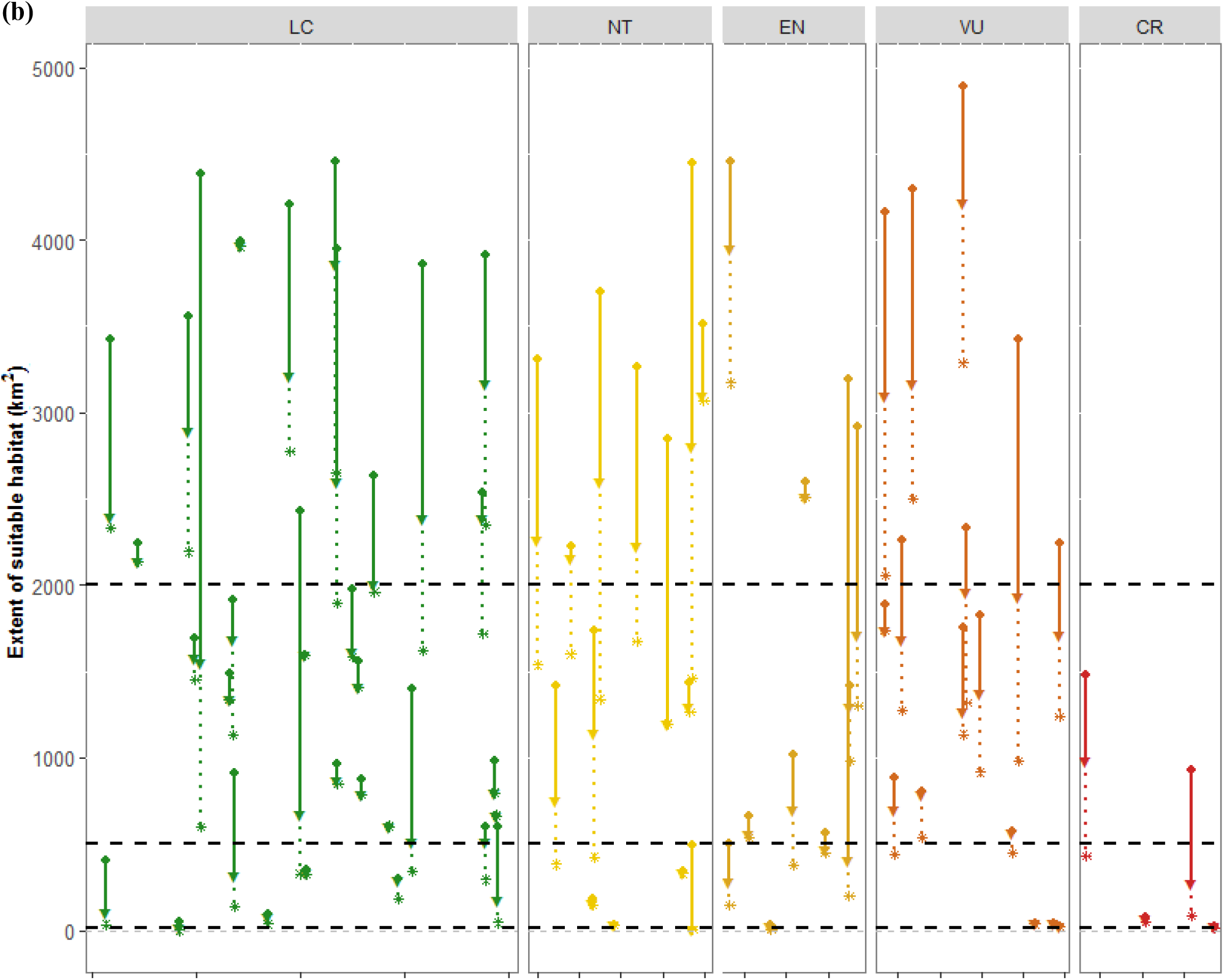
**(a)** Loss of extent of potential habitat for forest-dependent Colombian birds. Change in suitable habitat for each of the 550 species, until 2015 and the projected loss by 2040, split by each species’ current IUCN Red-list status: critically endangered (CR), endangered (EN), vulnerable (VU), near threatened (NT), and least concern (LC). The circles represent the historical extent of suitable habitat within the range of each species, the triangles the extent in 2015 and the asterisks the projected extent for 2040; the lines are drawn between the circle and triangle for the same species to highlight the species-specific change, the same for the dashed lines between the circles and asterisks. **(b)** The same figure but only showing species with historical extent of suitable habitat smaller than 5000 km^2^. In this graph the horizontal black dotted lines represent the thresholds for classification as vulnerable (VU) (2000 km^2^), endangered (EN) (500 km^2^) and critically endangered (CR) (10 km^2^) based on the IUCN criterion B2.

### Deforestation impact on assemblages

The impact of deforestation on forest-dependent bird assemblages in Colombia was dramatic. The Loss Index for the country was 35 (35% of Colombia’s forest-dependent species had lost at least 35% of their suitable habitat by 2015) (see Supplementary Table 1 and 2 for full details). Using the land use change model, the LI was projected to increase to 43 by 2040. Moreover, 18% of the species (n=99) had lost more than half of their historical habitat in the country by 2015 and 38% are projected to lose 50% or more of their habitat by 2040 (Fig. 2).

**Figure 2.**
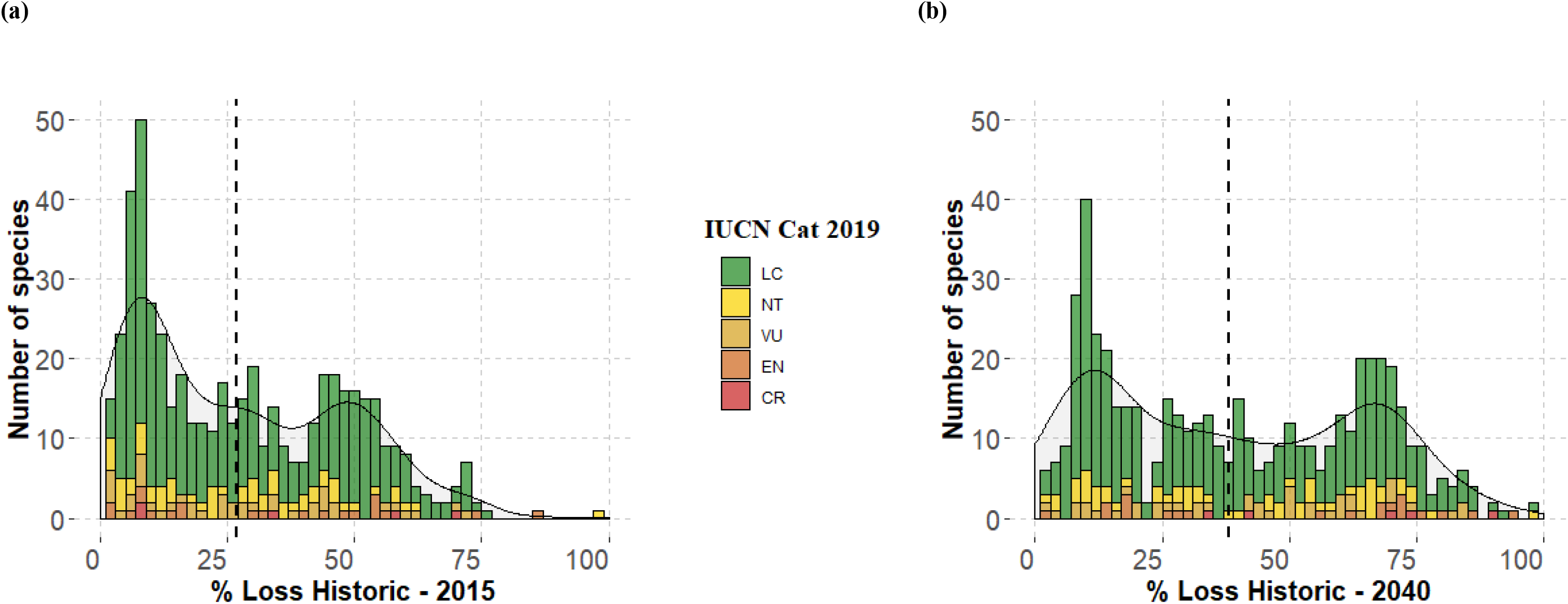
**(a)** Histogram of the number of forest-dependent birds (n=550 species) with different proportions of historical habitat loss until 2015 and **(b)** the projected for 2040 in Colombia. Continues lines represent a smoothed count estimate, vertical dashed lines show the mean habitat loss.

There were large differences in 2015 LI values for regions that differed in land-use intensity and biogeographic characteristics (Fig. 3a). For example, in the Caribe region, the LI was 82. This high LI value showed that forest loss had far-reaching effects on the majority of this region’s forest avifauna, despite the fact that only 15 species occurring in the region are listed as threatened. The LI dropped to 28 and 14 for the Amazon and Pacific regions, respectively, where forest loss has not been as extensive (Fig. 3a). However, in the species rich Amazon foothills, each km^2^ of forest loss has the potential to affect up to 230 forest-dependent species (Fig. 3a-b).

**Figure 3.**
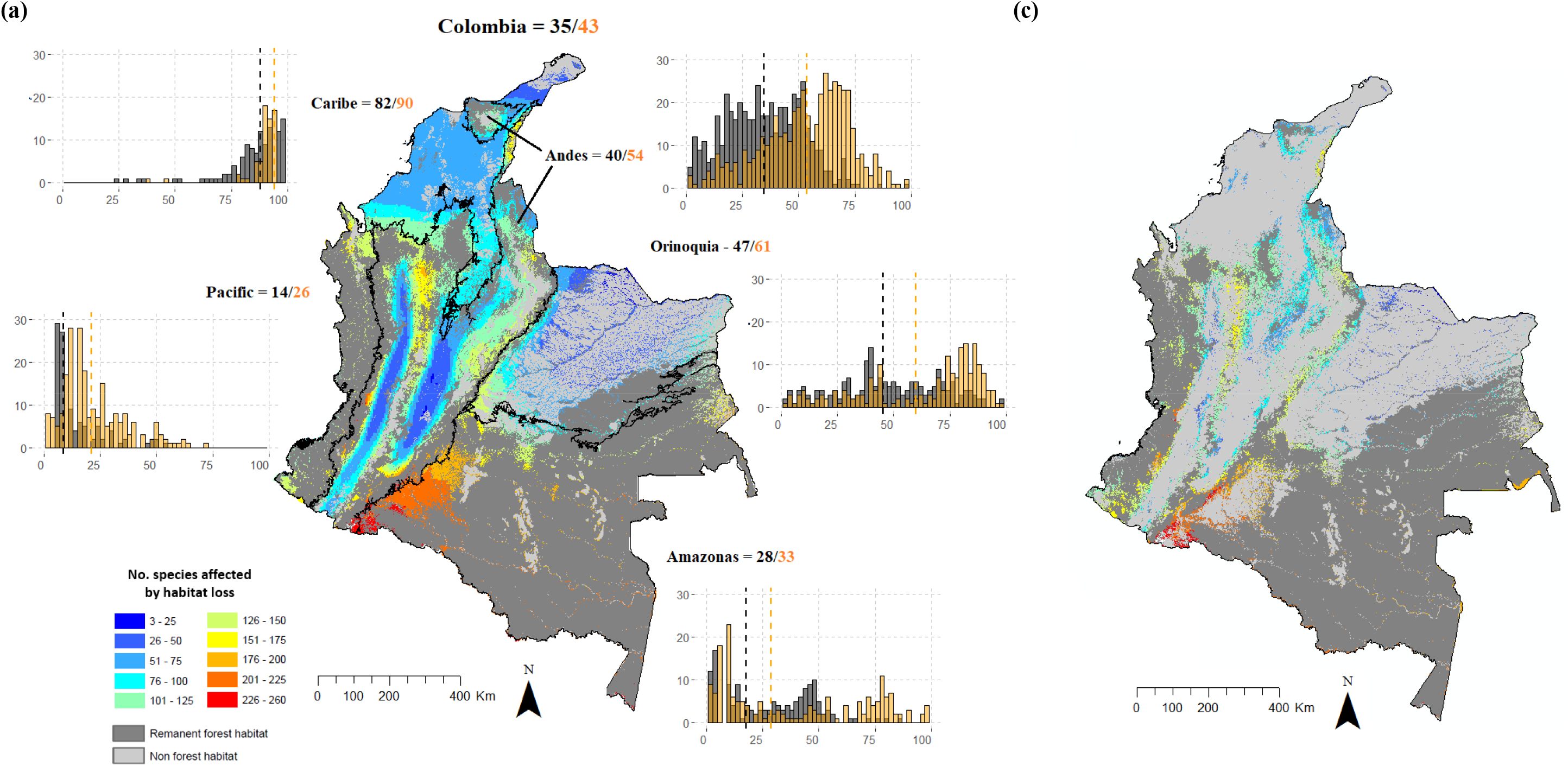
**(a)** Habitat loss index (LI) and histogram of the number of forest-dependent birds against percentage of historical habitat loss until 2015 (black) and the projected for 2040 (orange) for Colombia and its different biotic regions. Vertical dashed lines in the histograms show the mean habitat loss. In the centre a map of the number of forest-dependent birds potentially affected by habitat loss in different regions of Colombia (dark grey; remnant forest habitat, light grey; non forest habitat; colour key; number of species that potentially lost natural habitat) to 2015 **(b)** and from 2015 to 2040. Caribe (n=170 species); Andes (n=491 species); Orinoquía (n=213 species); Amazonas (n=301 species); Pacific (n=285 species).

The Andes region, which had the most forest-dependent species (491 species), had the greatest projected increase in its LI, from 40 in 2015 to 54 in 2040 (Fig 3a). The region with the highest LI by 2040 was Caribe with 90. The Amazon and the Pacific regions maintained relatively low LI, 33 and 26 respectively, However the LI for the Pacific region almost doubled from the value in 2015 (LI = 14).

For regionally-endemic species, the LI was 43 in 2015 and is projected to be 53 by 2040 (Fig. 4a) (see Supplementary Table 1 for full details). This implies that more than half of Colombia’s regionally-endemic species are projected to lose at least half of their habitat by 2040. The areas where deforestation is projected to affect more regionally-endemic species is the north-east of the Antioquia department where deforestation has already affected the habitat of more than 15 regionally-endemic species (Fig. 4b-c).

**Figure 4.**
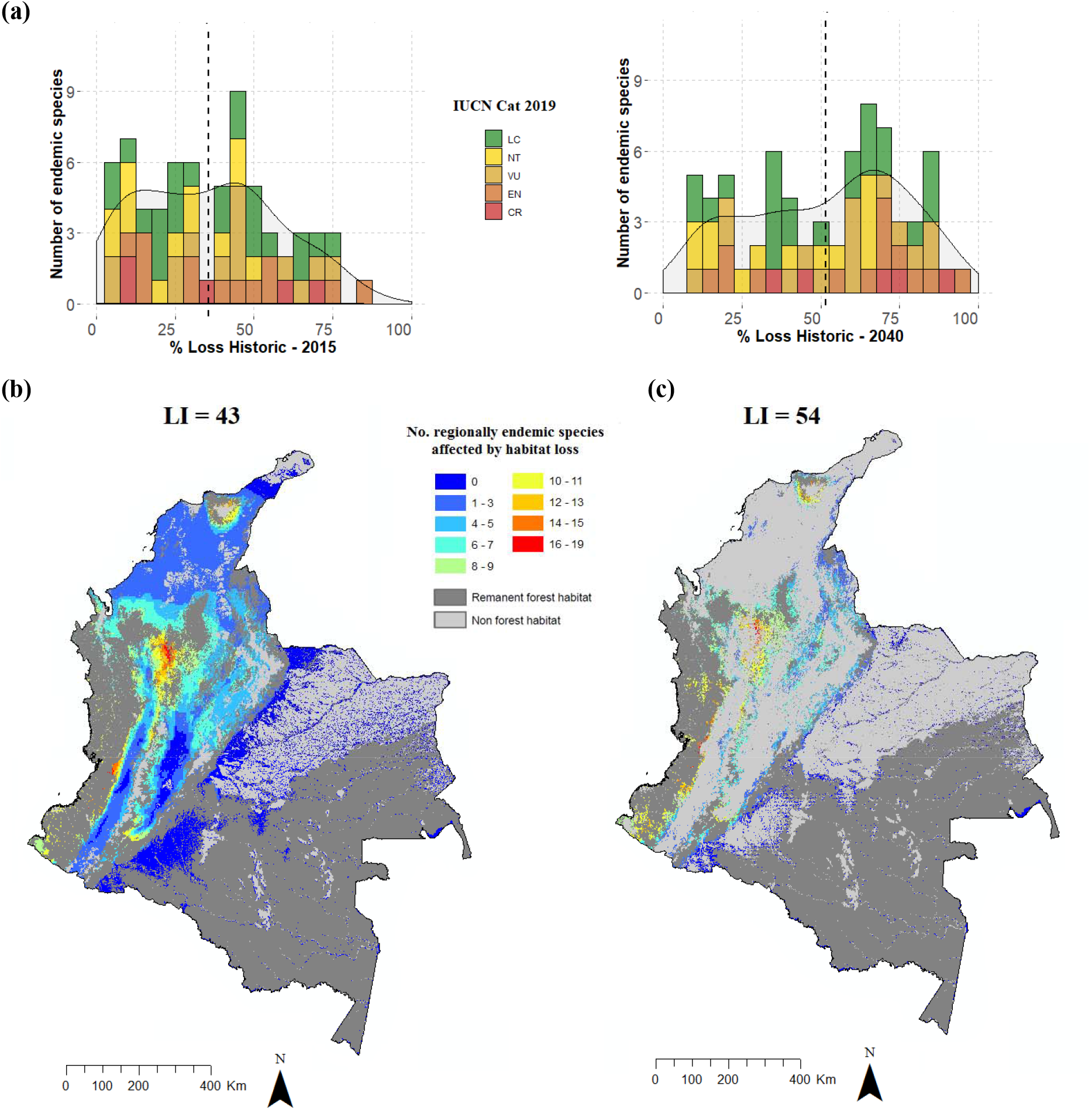
**(a)** Histogram of the number of regionally-endemic forest-dependent birds (n=69) against the percentage of historical habitat loss until 2105 and the projected for 2040 in Colombia. The vertical black dashed lines show the mean habitat loss. Continuous lines represent a smoothed count estimate. **(b)** Habitat loss index (LI) and map of the number of regionally-endemic forest-dependent birds potentially affected by habitat loss in different regions of Colombia (dark grey; remnant forest habitat, light grey; non-forest habitat; colour key; number of species that potentially lost natural habitat) for 2015 and **(c)** 2040.

We also explored the LI for four of the largest species groups of Colombia’s forest-dependent bird assemblages. Tanagers, which based on our assessment are the most species rich group of forest-dependent species (61 species), had a LI of 43 in 2015, which increased to 51 when projected forest loss by 2040 was included (Supplementary Fig. 2a). Ovenbirds (n=47) had a LI of 38 in 2015 with a projected increase to 42 by 2040 (Supplementary Fig. 2b); flycatchers (n=44) had a LI of 36 in 2015 projected to increase to 44 by 2040 (Supplementary Fig. 2c); and antbirds (n=53) had a LI of 27 in 2015 projected to increase to 32 by 2040 (Supplementary Fig. 2d).

### Deforestation impact on individual species

Deforestation has been and is projected to have major impacts on several threatened and non-threatened species in Colombia (Fig. 1 & 2) with 40% (n=208) of all forest-dependent species projected to lose at least half of their habitat by 2040, and 15% (n=79) projected to lose 70% or more of their habitat by 2040. The maximum proportional loss of habitat to 2015 and to 2040 among all the forest-dependent species was for the Giant Conebill *(Conirostrum binghami)* with 99%. This species was also the one with the maximum proportional loss between 2000 and 2015 (recent loss) with 63% of the remaining habitat to 2000 lost in this period. The maximum projected proportional loss between 2015 and 2040 was for the Western Hemispingus *(Sphenopsis ochracea)* with 93% of its remaining habitat in 2015 projected to be lost in the period to 2040 (see Supplementary Table 2 for full details).

The maximum proportional loss of historical habitat for regional endemics to 2015 and to 2040 was for the Antioquia Wren *(Thryophilus sernai)*, with 87% and 94% respectively. The maximum proportional loss between 2000 and 2015 (recent loss) was for the Tolima Dove (*Leptotila conoveri*), with 31% of the remaining habitat in 2000 lost in this period, and the maximum proportional projected loss between 2015 and 2040 was for the Chestnut-capped Piha *(Lipaugus weberi)* with 69% of the remaining habitat in 2015 projected to be lost in this period (Fig. 5; see Supplementary Table 1 for full details).

**Figure 5.**
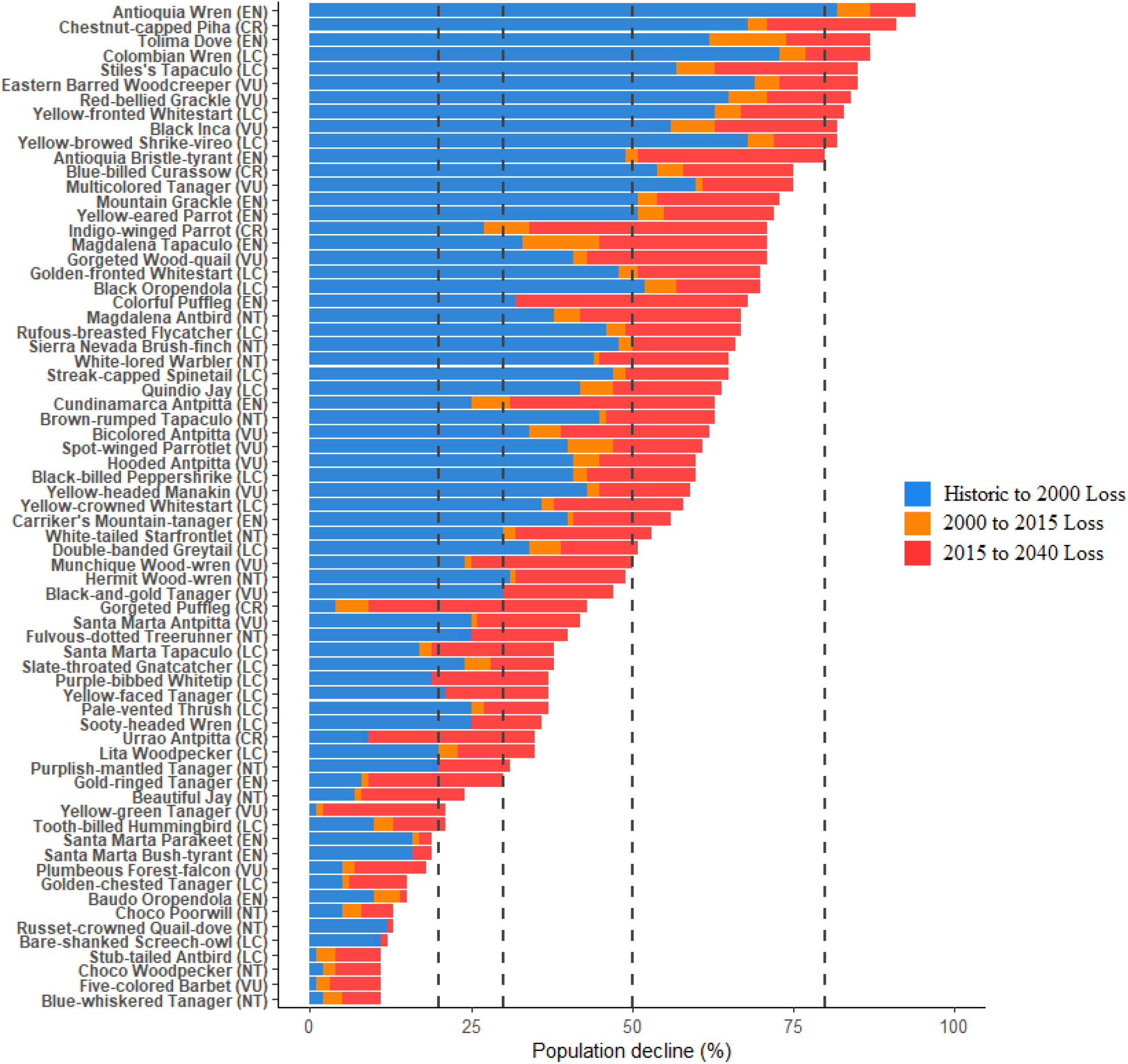
Population declines from habitat loss. The blue bar is the habitat loss until 2000, the orange bar the habitat loss until 2015 and the red bar the projected habitat loss by 2040. This graph only shows the 69 regionally-endemic forest-dependent species. Vertical lines represent the thresholds for classification as near threatened (NT) (20%), vulnerable (VU) (30%), endangered (EN) (50%), and critically endangered (CR) (80%) based on the IUCN criterion A4. In parenthesis IUCN category in 2019.

### Threat assessment for regionally-endemic species

Our threat assessment for the 69 regionally-endemic species showed that 49 species lost 20% or more of their historical habitat by 2015, 40 species lost 30% or more, 18 species lost 50% or more and one species lost more than 80%. When the projected loss until 2040 is included, 57 species are projected to lose 20% or more of their historical habitat by 2015, 54 species are projected to lose 30% or more, 39 species are projected to lose 50% or more and 11 species are projected to lose more than 80% (Fig. 5). Of the 18 species that had lost 50% or more of their historical habitat by 2015, six are not currently classified as threatened by the IUCN, and of the 39 species that are projected to lose 50% or more of their historical habitat by 2040, 12 are not currently classified as threatened by the IUCN (see Supplementary Table 3 for full details).

When considering recent loss only (2000 – 2015; i.e. not loss compared to historical coverage) our results show that only two species lost more than 20% of their habitat. However, when the projected loss between 2015 and 2040 is included in this assessment (2000 – 2040) 49 species are projected to lose 20% or more of their historical habitat by 2040, 37 species are projected to lose 30% or more and 15 species are projected to lose more than 50% (Supplementary Fig. 3). Of the 15 species that are projected to lose, by 2040, 50% or more of their remaining habitat in 2000, three are not currently classified as threatened by the IUCN (see Supplementary Table 3 for full details).

Twenty-four species had less than 3000 km^2^ of area of occupancy by 2015, 17 species had less than 2000 km^2^ and seven had less than 500 km^2^. When this assessment is done based on the projected extent of area of occupancy by 2040, 29 species had less than 3000 km^2^ of area of occupancy by 2040, 24 species had less than 2000 km^2^, 10 had less than 500 km^2^ and 1 had less than 10 km^2^ (see Supplementary Table 3 for full details). Of the 24 species that are projected to occupy less than 2000 km^2^ by 2040, four are not currently classified as threatened by the IUCN (see Supplementary Table 3 for full details).

## Discussion

Our study highlights the importance of assessing the impact of deforestation on whole species assemblages. We report a Loss Index (LI) of 35 for Colombia in 2015: 35% of the species analysed had lost 35% or more of their historical habitat. If recent deforestation trends continue, we estimate that LI will increase to 43 by 2040. Despite this ongoing attrition of habitat, only 17% (n=35) of the species that are projected to lose 50% or more of their habitat in Colombia by 2040 (n=208) are currently classified as threatened by the IUCN (2019b), suggesting that there are many species that are facing an imminent extinction threat from habitat loss even though they are not formally listed as threatened.

### Depletion of Colombia’s forest bird habitat

Deforestation has impacted almost all the forest-dependent bird species in the country (n=536; 96.5%) but most still retain more than half of their historical habitat (n=451, 82%). However, if deforestation trajectories continue, 40% (n=208) of the forest-dependent species would lose half or more of their suitable habitat in the country by 2040 (Fig. 2). This is particularly concerning for currently non-threatened species which comprise 83% of the assemblage we explored, and which normally are not the focus of conservation initiatives.

When assessing only regionally-endemic species, our results show a more dramatic change than the one observed for the entire forest-dependent bird assemblage. Seventy-four percent (n=51) of the species retained at least half of their historical habitat in 2015, but if deforestation trajectories continue, 57% (n=39) of the species would lose half or more of their historical habitat by 2040 (Fig. 4a).

Our results also showed that forest-dependent tanagers were the bird family with the greatest proportional loss of suitable habitat in the country to 2015 and projected to 2040. Tanagers are particularly diverse in the Andes, which is one of the most heavily historically altered regions of Colombia (Etter & van Wyngaarden 2000; Etter et al. 2006a). More knowledge of the threats imposed by deforestation to tanager habitat in the Andes is needed in order to avoid their extinction.

### Regional variation in impact of deforestation on assemblages

At a regional scale our results show that the Caribe region had the highest Loss Index value (LI = 81) and had a mean historical habitat loss of 88% by 2015. This reflects high levels of forest loss that is affecting most of the species in the assemblages in this region. Past studies have also identified the Caribe as a priority region due to its high land conversion and low protection levels (Forero-Medina & Joppa 2010; Negret et al. 2020), highlighting the importance of conservation aimed at reducing forest loss in this region. Colombia’s relatively well-preserved Choco and Amazon regions had low LI values and the mean loss of historical habitat was below 20% for both. These regions are characterized by high species richness with Choco representing a hotspot for endemic birds (Mittermeier et al. 2004). The maintenance of intactness in these regions is critical as diverse and intact species assemblages underpin ecosystem functions such as seed dispersal, pollination, pest control and carbon sequestration (Cardinale et al. 2012; Watson et al. 2018; Maxwell et al. 2019). The Andes region, which was the most bird species rich, was also the region with a more drastic projected change in the LI (from 40 to 54) and in mean habitat loss (35 to 54) when compared to the other regions (Fig. 3a). Actions focused on preventing the loss of habitat for endangered and range restricted species in this region are needed.

The west section of the Amazon foothills includes areas of very high species richness (overlapping ranges of >200 forest-dependent birds in places) (Fig. 3). This region has undergone a rapid increase in deforestation since the peace deal (Clerici et al. 2018; Murillo Sandoval et al. 2020), so investment in management of protected areas and other effective area-based conservation measures is a priority to avoid the loss of these rich bird assemblages. Our results also showed that the central and eastern Andes are areas where deforestation has substantially affected forest-dependent tanager species, while forest-dependent antbirds were especially affected in the Amazon-Andes foothills in Putumayo. Tanagers and antbirds provide important ecosystem services such as seed dispersal, pollination and insect population regulation since their diet is mostly fruits and insects (Hilty & Brown 1986; McMullan et al. 2010), making their effective conservation crucial for ecosystem functioning and ecosystem service provision. Additionally, the north-east of the Antioquia department is an area where projected deforestation will affect bird assemblages that have a particularly high concentration of endemic species (Fig. 4). This region has been considerably affected by armed conflict, but there are several conservation initiatives and research expeditions in this area after the peace deal (United Nations 2018). These conservation initiatives would help protect the endemic bird species in this region.

### Implications for species threat status

Half of the regionally-endemic species are currently not classified as threatened based on the IUCN threatened species status criteria (see Supplementary Table 1 for full details) but our analysis suggests that many of those species are facing significant habitat loss that could lead to their extinction (Fig 5, Supplementary Fig 3). For example the Colombian Wren (*Pheugopedius columbianus*) and the Stiles’ Tapaculo (*Scytalopus stilesi*), which are currently classified as Least Concern (LC), have already lost more than 60% of their habitat in Colombia and are projected to lose more than 85% by 2040 (Fig 4). Targeted research for these and other species with similar characteristics is needed in order to identify the factors that are generating the reduction of their habitat and to determine if a revision of their Red List classification is warranted. In contrast, loss of suitable habitat has been small for some threatened and near threatened species in Colombia. This in part is associated to the fact that some of these species inhabit less accessible areas (e.g., Wattled Curassow [*Crax globulosa*] and Banded Ground-cuckoo [*Neomorphus radiolosus*]) or that have been the target of successful conservation actions (e.g., Urrao Antpitta [*Grallaria fenwickorum*] and Gorgeted Puffleg [*Eriocnemis isabellae*]) (Carantón-Ayala & Certuche-Cubillos 2010; ProAves 2019). Moreover, our threat assessment for regionally-endemic species showed that accounting for projected habitat loss can provide useful information of the species that need proactive conservation interventions as projected loss would affect their habitat, population numbers and threat status.

### Limitations of our analysis

While our method is a rapid way to assess and describe the impacts of deforestation to the habitat of forest-dependent species, some limitations have to be noted. In predicting extent of and change in potentially suitable habitat, the underlying maps, while the most comprehensive and detailed up to date, likely contain commission and omission errors which can lead to inaccurate forest change estimates (Rondinini et al. 2011; Ficetola et al. 2014; Tracewski et al. 2016; Palacio et al. 2020). For some species in heavily historically altered regions, like the Caribe (Etter & van Wyngaarden 2000; Etter et al. 2006a), mapped ranges might be more consistent with the species’ current area of occupancy, reflecting the remnants of the species pre-human range, and thus not account for range contractions which may have occurred and been caused by historical habitat loss (e.g, Blue-billed Curassow [*Crax alberti*]). Despite this, species in the tropics tend to have conservative niches, making geographical and climatic barriers more effective (Janzen 1967; Brown 2014). This in turn makes species distribution change less common. Additionally, species for which habitat reduction represents the main extinction risk are normally those that are restricted to smaller areas and have more specialized niches (Harris & Pimm 2008; Birand et al. 2012) so their historical distributions can be better inferred. Even though this should be the case for the majority of species, we are aware that there are information gaps in relation to the bird communities present in some regions of the country, and that there are constant reports of records beyond the known range and altitudinal limits for several species (Negret & Laverde-R. 2014; Gomez-Bernal et al. 2015; Negret et al. 2015). Based on this we decided not to generate further altitudinal range refinements for the species analysed, making our estimates of the impact of deforestation to the habitat and population size of forest-dependent species conservative.

There is also a lack of understanding of habitat preferences for some species (e.g., Magdalena Tapaculo [*Scytalopus rodriguezi*]), especially species with highly specialised habitat requirements or that exist at very low densities within suitable habitat. For example, based on our results the Banded Ground-cuckoo (*Neomorphus radiolosus*), which is categorized as Endangered (EN), has retained more than half of its habitat. However, the extent of suitable habitat for the species might be lower than our estimates suggest as it is a species present in low densities, and for which ecological characteristics are poorly understood (Hilty & Brown 1986; McMullan et al. 2010; Ayerbe-Quiñones 2018). This is because our broad classification of potential ‘habitat’ – forest – does not allow us to account for species-specific specialisation (although this is accounted for broadly by restricting habitat amount calculations to within the current range of where species occur). Consequently, while our results represent an improvement on existing knowledge in the majority of cases, each assessment must be judged carefully and in combination with on the ground information if possible. Finally, our results should be used in context with each species’ particular circumstances, considering other threats such as the impacts of selective logging (Isaac & Cowlishaw 2004; Mayor et al. 2015), hunting (Isaac & Cowlishaw 2004; Benítez-López et al. 2017), and wildlife trade (Symes et al. 2018) which will likely cause substantial reductions in some species, as would occur in any normal IUCN assessment process.

## Conclusion

Our results show the importance of holistic assessments of the impact of deforestation on whole species assemblages. We show the level of historical and recent reduction in habitat for non-threatened forest-dependent species in Colombia is comparable to that of threatened species and we argue that monitoring of both threatened and non-threatened species must be done in tandem. Moreover, many of the non-threatened species that have already lost half or more of their potential habitat in Colombia are considered ‘common’ (e.g., Basileuterus tristriatus[*Three-striped Warbler*]) or ‘abundant’ (e.g., Grey-breasted Wood-wren [*Henicorhina leucophrys*]) (Hilty & Brown 1986; McMullan et al. 2010). Common and abundant species represent the bulk of individuals in forest assemblages and their ecological roles are fundamental for ecosystem functioning (Sekercioglu 2006; Gaston & Fuller 2008) and the provision of ecosystem services (Sekercioglu 2012; Gaston et al. 2018). Their extirpation can reduce income for local communities in ecotourism (Maldonado et al. 2018), reduce pest control in nearby agroforest and agricultural landscapes (Sekercioglu 2012) and reduce pollination and seed dispersal of fruiting and timber trees in adjacent agricultural landscapes (Gaston & Fuller 2008).

While identifying and halting the underlying causes of deforestation is fundamental to avoid the extinction of forest-dependent species, improving the protection for species under imminent risk is necessary in the short term to avoid their extinction. Understanding future threats to species based on projected impacts of deforestation is therefore an important tool which helps to generate better conservation plans and actions to protect focal species of otherwise unseen risk. We believe the methodological framework applied in this study can provide a rapid way to generate an initial evaluation of the state of entire species assemblages in other biodiverse countries and regions where field data is not complete or available.

## Acknowledgements

We are grateful to R.D. Palacio, J.R. Allan and K.R. Jones, for providing constructive feedback and discussion around elements of this study.

## Authors’ contributions

P.J.N. conceived and designed the study with J.S.S., M.M. and J.E.M.W suggestions. P.J.N. implemented the analysis. All the authors contributed to subsequent drafts and gave final approval for publication.

## Data Availability

Supplementary Table 1,2 & 3 will be made available upon request, previous to its deposition in an open-access repository with the peer-reviewed version of this study. Requests should be sent to the corresponding author.

## Supplementary material

Supplementary Table 1, 2 & 3 are in an excel file

**Supplementary Figure 1.**
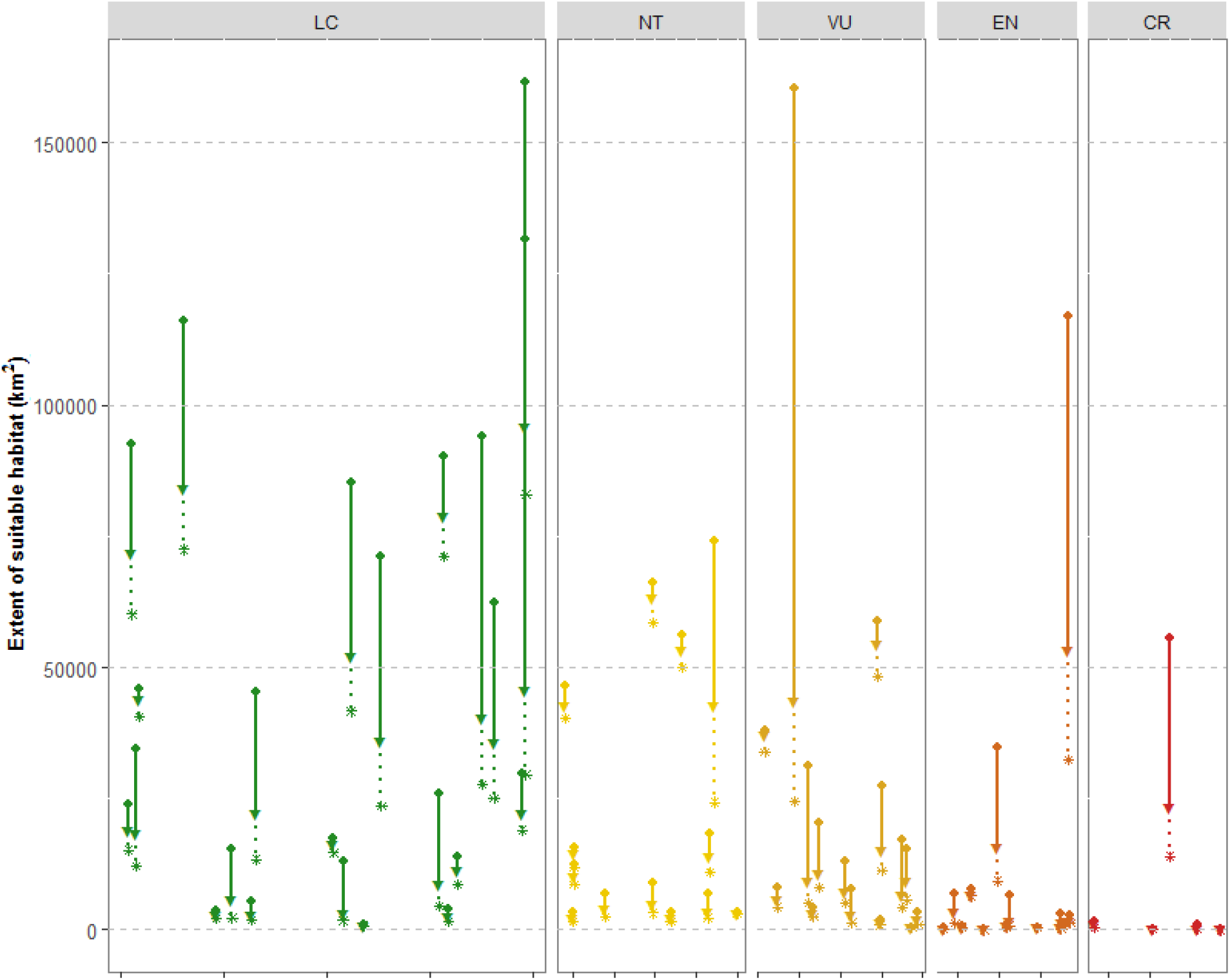
**(a)** Loss of extent of suitable habitat in Colombia for regionally-endemic forest-dependent birds. Change in suitable habitat for each of the 69 species until 2040, split by their current IUCN Red-list status: critically endangered (CR), endangered (EN), vulnerable (VU), near threatened (NT), and least concern (LC). The circles represent the historical extent of suitable habitat within the range of each species, the triangles the extent in 2015 and the asterisks the projected extent for 2040; the lines are drawn between the circle and triangle for the same species to highlight the species-specific change, the same for the dashed lines between the circles and asterisks.

**Supplementary Figure 2.**
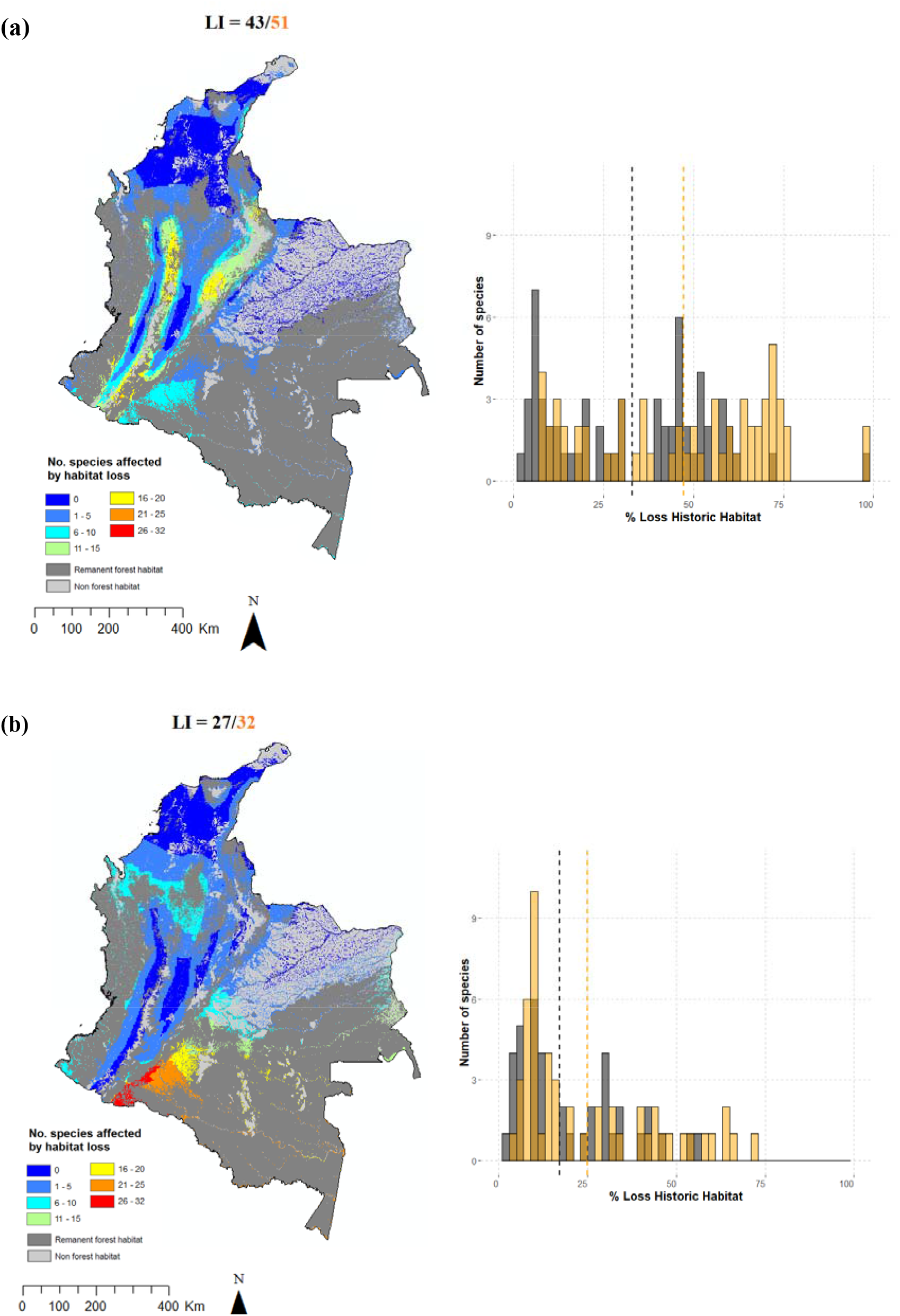
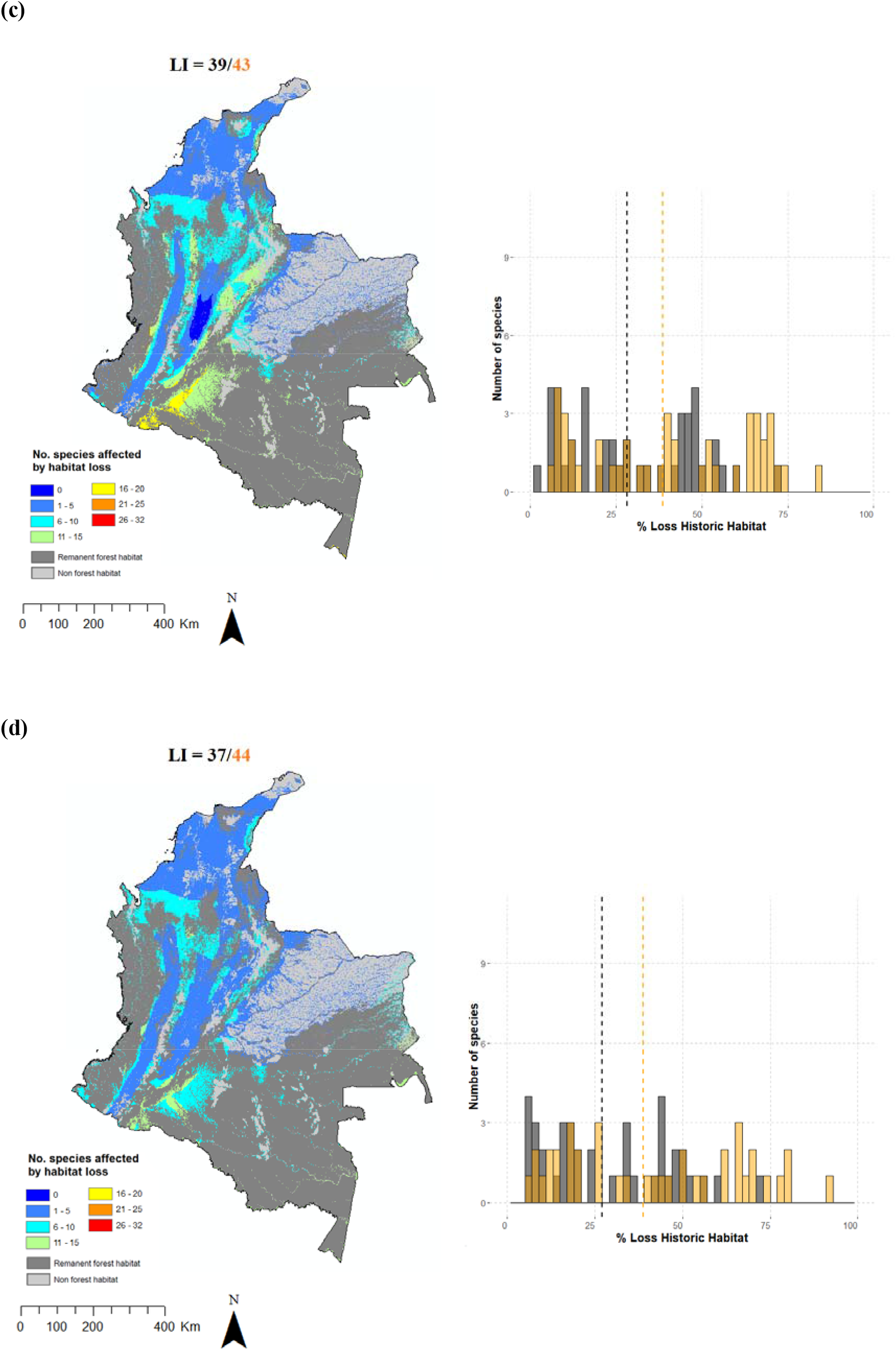
Habitat loss index values (LI), map of the locations of potential lost natural habitat (shaded pixels), and histogram of the number of forest-dependent species against percentage of historical habitat loss until 2105 (black) and the projected for 2040 (orange) in Colombia for 4 major bird groups. Dashed lines show the mean habitat loss by 2015 and the projected by 2040 **(a)** Tanagers (Thraupidae, n = 61 species), **(b)** Antbirds (Thamnophilidae, n = 53 species), **(c)** Ovenbirds (Furnariidae, n = 47 species) and **(d)** Flycatchers (Tyrannidae, n = 43 species). In the maps, dark grey depicts remanent forest habitat, light grey non forest habitat and the colour key, the number of species that potentially lost natural habitat.

**Supplementary Figure 3.**
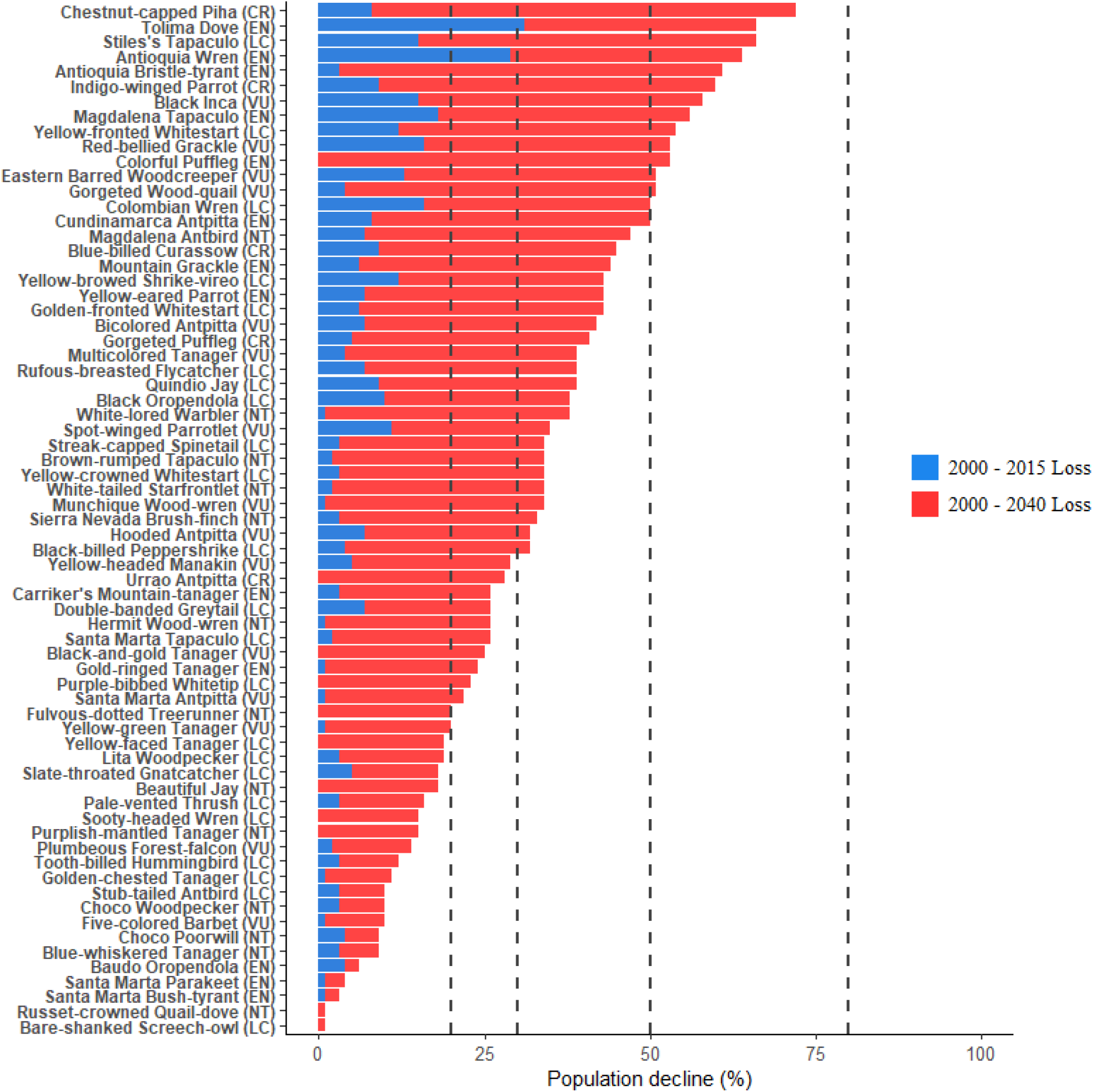
Population declines from habitat loss. The blue bar is the habitat loss from 2000 to 2015, and the red bar the projected habitat loss by 2040. This graph only shows the 69 regionally-endemic forest-dependent species. Vertical lines represent the thresholds for classification as near threatened (NT) (20%), vulnerable (VU) (30%), endangered (EN) (50%), and critically endangered (CR) (80%) based on the IUCN criterion A4. In parenthesis IUCN category in 2019.

